# Rapid population recovery of a foundation species from experimental small-scale harvesting

**DOI:** 10.1101/2023.12.14.571731

**Authors:** Eliseo Fica-Rojas, Daniela N. López, Alejandro Pérez-Matus, Nelson Valdivia

## Abstract

Ecological stability is central to understand how disturbances challenge the persistence of populations and communities through time, particularly when species with strong effects on other species are disturbed. The bull kelp *Durvillaea incurvata* is a foundation habitat forming species that provides habitat, food, and shelter for various species, and supports the livelihoods of human communities along the southeast Pacific coast of Chile. Harvesting of *D. incurvata* has raised concerns about the long-term viability of its populations, but the stability responses of bull kelps to anthropogenic disturbances are still unclear. Here, we conducted a manipulative field experiment in which we removed once all individuals of the bull kelp from two sites in southern Chile. We simulated the loss of bull kelps to harvesting in 1-m^2^-plots interspersed in matrices of dense *D. incurvata* stands. Fronds cannot regrow from the holdfasts after harvesting. Holdfasts were therefore also removed, a practice not typically carried out by local gatherers. For 25 months we quantified bull kelp recruitment, holdfast densities, percent cover, mean frond size and density, biomass, and population size structure in two rocky intertidal sites. In both sites, all metrics completely recovered within five to seven months. The removal of *D. incurvata* did not have a significant impact on recruitment, which was constant during the experiment. The relatively small spatial scale of the disturbances, constant recruitment provided by the surrounding bull kelp matrix, and the removal of holdfasts that released settlement substratum may have allowed for the strong stability responses in these populations. Therefore, spatial heterogeneity, which allows for settles to colonize the disturbed plots, should be prioritized in management strategies of natural seaweeds stands.

## INTRODUCTION

Disturbances play a key role in structuring populations, communities, and ecosystems (Battisti et al., 2016; Newman, 2019). Both, anthropogenic and natural disturbances can induce direct mortality of organisms, modify resource availability, and induce changes in the temporal dynamics of aggregated functions such as community biomass or productivity (Battisti et al., 2016; Newman, 2019). These changes may extend to several spatial and temporal scales (Kéfi et al., 2019) and represent important societal concerns, due to the risk of losing ecosystem goods and services (Cardinale et al., 2012).

Ecological stability has been central to understand how natural systems respond to disturbances (Ives & Carpenter, 2007; Donohue et al., 2013; Kéfi et al., 2019). This property corresponds to the ability of populations and communities to resist and recover from disturbances, remaining relatively unchanged over time (Harrison, 1979; Pimm, 1984; Doak et al., 1998; Hillebrand et al., 2018). Stability encompasses multiple dimensions (or metrics) that allow us to describe different aspect of a response to the disturbance, such as resistance, resilience, and recovery, among others (Donohue et al., 2016; Kéfi et al., 2019; Hillebrand et al., 2018). Upon a disturbance takes place, resistance is the “immediate or early” response of a system to the disturbance, resilience is the speed or rate of return of the system to its dynamic reference state (Holling’s engineering resilience; Pimm 1984), and recovery indicates the state (relative to a reference state) at the end of an observational time (Donohue et al., 2016, Hillebrand et al., 2018).

Upon a disturbance takes place, population resilience and recovery can be estimated for several attributes that can be linked to aggregated properties such as population biomass—these attributes include, but are not limited to, individual sizes, densities, and percent cover of primary space, among others (Gladstone-gallagher et al., 2019; Kéfi et al., 2019, Valdivia et al. 2023). Studies usually have been focused on the understanding of single dimensions of stability or population attributes (Donohue et al., 2016, Kéfi et al. 2019). However, recent evidence suggests that single dimensions cannot reflect completely the overall stability of the system (Donohue et al., 2013, 2016; Hillebrand et al., 2018; Yang et al., 2019; Polazzo & Rico 2021). Also, estimating stability responses across several population attributes allows us to improve our understanding of how the effect of disturbances on demographic traits channel to aggregated population-level properties and functions that are relevant for management and conservation (i.e. population biomass; Cardinale et al., 2012; Gladstone-Gallagher et al., 2019).

Foundation species such as large laminarian and fucoid seaweeds (hereafter referred as kelps) are key within coastal ecosystems (Steneck, 2002; Steneck & Johnson, 2014; Cuba et al., 2022;). The large individual sizes and high densities of kelps contribute with high levels of biomass, sustaining coastal food webs and several ecosystem services (Graham et al., 2007; Aguilera et al., 2018; Edwards et al., 2020; Cuba et al. 2022). Also, canopies formed by kelp can modify local hydrodynamics and reduce environmental variability, regulating habitat availability and quality for other species in the understory and for early-stage conspecifics as well (Santelices, 1980; Taylor & Schiel, 2005; Toohey, 2006; Stachowicz et al., 2007; Steneck et al., 2008; Tait & Schiel, 2011). Therefore, assessing the population stability of foundation species is a key step to understand how coastal ecosystems interact with disturbances.

The bull kelp *Durvillaea incurvata* (hereafter referred to as *Durvillaea*) is one of the most characteristic foundation species along the temperate coast of Chile, where is intensely harvested for human consumption and for the alginate industry (Vásquez, 2008; Camus et al., 2019; Velásquez et al., 2020). This species is distributed between 39°S to 56°S in wave-exposed low intertidal and shallow subtidal rocky shores, covering extensive areas and reaching high individual density (up to 80 % of the shore and up to 30 individuals per m^2^; Santelices et al., 1980; Westermeier et al., 1994). The thallus of a *Durvillaea* individual is composed of a holdfast that is attached to the rocky substratum, a large buoyant frond which is responsible for the photosynthesis, and a flexible and strong stipe that connects the holdfast and fronds (Santelices et al., 1980; Taylor & Schiel, 2005).

*Durvillaea* individuals show high growing rates (Santelices et al., 1980), can reach up to 10 meters long, live up to 10 years, and provide large amounts of biomass to the local community (up to 80 kg m^-2^; Santelices et al., 1980, Taylor & Schiel 2005). Individuals of *Durvillaea* are reproductive most of the year and females can produce millions of eggs that can, once are fertilized, settle on primary substrate within few hours (Taylor & Schiel, 2005) and produce massive recruitment episodes during warmer seasons (Westermeier et al., 1994). Often, some individuals are lost from the rocky substrate due disturbances occurring at short time intervals, such as storms (Santelices, 1980; Moreno, 2001; Taylor & Schiel, 2003), marine heat waves (Thomsen et al., 2019), and direct human harvesting (Moreno et al., 1984; Bustamante & Castilla, 1990; Parra et al., 1992; Castilla et al., 2007). However, rapid generation times, high reproductive output, and fast-growing rates of *Durvillaea* (Santelices et al., 1980) should promote population resilience and complete recovery within one or few years. Thus, following a disturbance, we should expect several attributes of these population to recover within a short period of time (i.e., within a year).

On the other hand, high densities of large holdfasts and fronds reduce environmental variability and enhance recruitment, survival, and growing of early stages of conspecifics (Ojeda & Santelices, 1984; Taylor & Schiel, 2003, 2005; Castilla et al., 2007; Layton et al. 2019), promoting self-replacement after disturbances. In addition, local harvested usually remove only the fronds from each individual and leave the holdfast. Since fronds cannot regrow from these holdfasts, these structures reduce the cover of substratum that would be otherwise available for settlement of bull kelp propagules (Westermeier et al., 1994). Thus, the loss of kelps may disrupt post settlement survival, recruitment, and individual growth, which may lead to a reduced resilience and recovery of several populations’ attributes in the short temporal scale (within few years).

In this study, we conducted a field-based manipulative experiment in two sites and for 25 months to assess the resilience and recovery of several attributes of *Durvillaea* populations impacted by a pulse disturbance that removed all the individuals from experimental plots and that simulates the magnitude by which gatherers harvest *Durvillaea* population in south-central Chile. We focused on attributes associated to individuals’ densities, percent cover, frond sizes, population biomass, and size structure. We tested two competing hypotheses:

**Hypothesis 1**: Since *Durvillaea* has a high reproductive output, massive recruitment, and high growing rates (Santelices et al., 1980; Taylor & Schiel, 2005), we predict fast resilience and high recovery of population attributes and biomass within a year or few months after the removal of kelps.

**Hypothesis 2**: In contrast, and since the high density of *Durvillaea’* large individuals improve population recruitment and post settlement survival and growing (Taylor & Schiel, 2005; Castilla et al., 2007), we predict that kelp removal should decrease recruitment and lead to low resilience and an incomplete recovery of several population attributes at the intra-annual temporal scale.

## MATERIAL AND METHODS

### Study sites and experimental design

The study was conducted in south-central Chile (∼39° S), within an intermediate biogeographic province located between the Peruvian and Magellan coastal biogeographic provinces (Camus, 2001). Here, two intertidal rocky shores separated by approximately 40 km, Chaihuín (39°56′56″S; 73°34′25″W) and Loncoyén (39°49’19.2”S; 73°16’48.2”W), were used to carry out a field manipulative experiment. Chaihuín is under a Territorial Use Rights for Fishing (TURF) scheme for marine resources associated to *Durvillaea* (Carrillo et al. 2015), where locals manage species such as *Durvillaea,* limpets (*Fissurella* spp.), Locos (*Concholepas concholepas*), and others. Loncoyén is an open access area, where harvesting of many species takes place without any legal regulation. In both sites, direct harvesting of *Durvillaea* takes place mainly during low tides of spring and summer months.

In each site we deployed twenty 1-m^2^ plots marked with stainless-steel bolts and a numbered plastic plate. All plots were evenly assigned to five blocks randomly distributed along the shore (four plots per block). Each plot was haphazardly assigned to one of two treatments: either pulse disturbed or control. The pulse disturbance treatment consisted of a single disturbance that removed once all the individuals of the dominant kelp *Durvillaea* from the substrate. In this way, the disturbance constituted an actual perturbation (sensu Rykiel 1985), because the disturbance generated an evident change in *Durvillaea* population. Kelps were removed with aid of chisels and knifes. The control treatment consisted of plots in which kelp was not manipulated. We quantified six *Durvillaea* populations attributes (see *Statistical analyses*, below) before the disturbance was applied, two months after the disturbance, and then every 3 to 6 months. The experiment was initiated in January and October of 2020 and finalized in February and November 2022 in Chaihuín and Loncoyén respectively (i.e., 25 months of monitoring). All manipulations and observations were conducted during diurnal low tide hours.

### Morphological traits and length – biomass calibration curves

All the individuals removed at the beginning of the experiment were measured and weighted for the construction of length-biomass calibration curves. These curves allowed us to estimate individual and population biomass repetitively in experimental plots without doing a destructive sampling.

After the experimental removal, the removed individuals of *Durvillaea* were carried within few hours to the laboratory, where samples were immediately measured and weighted to the nearest 0.1 cm and 0.1 g, respectively. For each sampled individual, holdfast diameter (cm), stipe diameter (cm), and frond length (cm) were measured with a measuring tape. *Durvillaea* thalli commonly show several stipes and fronds growing from a single holdfast, because of coalescence of several embryos during settlement (Santelices et al., 1980) or from holdfast fusion of early-stage individuals (Castilla & Bustamante, 1989). Usually, coalescent individuals can contain several fronds of different sizes (up to 16 fronds per holdfast in our dataset). Therefore, for coalescent individuals, all each stipes and fronds were measured. After measuring, the wet weight of the whole holdfast, stipes, and fronds were estimated and then, all the kelp material was dried at 70° C for 48 h in a dry air oven. With these measures we developed the length-biomass relationships separately for holdfasts, stipes, and fronds of each site, using exponential regression models:

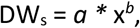

where DW is the dry weight (g), **s** is the kelp structure (holdfast, stipe, or frond), *a* is the intercept, x the length or diameter (cm) of kelp structures, and *b* is the slope of the exponential equation (see parameter estimates in Table 1).

**Table 1.** Exponential regression coefficients of biomass estimation for holdfast, stipes and fronds of *Durvillaea incurvata* individuals from two sites in southern Chile (Chaihuín and Loncoyén). HD: Holdfast diameter (cm), SD: Stipe diameter (cm), FL: Frond length (cm).

|  | Chaihuín |  |  | Loncoyén |  |  |
| --- | --- | --- | --- | --- | --- | --- |
| Kelp structure | Equation | r<sup>2</sup> | n | Equation | r<sup>2</sup> | n |
| Holdfast dry weight (HDw) | $0.212 * HW^{1.0496}$ | 0.9943 | 100 | $0.2412 * HW^{1.0522}$ | 0.9609 | 129 |
| Stipe dry weight (SDw) | $0.1486 * SW^{1.0599}$ | 0.9973 | 217 | $0.1749 * SW^{1.0382}$ | 0.9409 | 222 |
| Frond dry weight (FDw) | $0.1677 * FW^{1.0116}$ | 0.9933 | 87 | $0.1888 * FW^{1.0185}$ | 0.9260 | 158 |

### Population dynamics

For each plot, we quantified the temporal dynamics of population attributes associated to the use of primary space and those associated to the canopies formed by *Durvillaea*. For instance, the percent cover of *Durvillaea* within each plot was visually estimated, with aid of a 1-m^2^ and 1-% accuracy quadrat, as the proportion of the substrate covered by kelp holdfasts. We estimated the density of juvenile and adults’ holdfasts as the number of holdfasts with a diameter larger than 1 cm and 5 cm respectively (Santelices et al., 1980; Castilla et al., 2007). Individuals with a holdfast smaller than 1 cm in diameter were considered as “recruits”. When recruit densities were too high, we counted and measured the individuals in one quarter (0.25 m^2^) of the plot and then data were extrapolated to m^2^. Also, we measured the frond length (cm) of each individual found in experimental plots with a plastic measuring tape. In those cases of coalescent individuals, we considered all the fronds contained in a single holdfast as separate individuals. Then, we estimated the mean frond density and the mean frond length of each plot, considering clumped all juvenile and adult individuals within experimental plots.

Additionally, with the morphological measures of each individual we estimated biomass as described in the previous section (see Morphological traits and individual biomass estimation, above). Then, for each sampling time and experimental plot, we estimated population biomass as the summed biomass of all individuals.

To assess the temporal dynamics of size structure, we estimated Gini inequality coefficient (*G)* for data of frond biomass as:

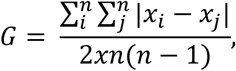

where x_i_ and x_j_ are the biomass of all possible pairs of individuals, x represents the mean biomass of the population, and n represents all the sampled individuals. This coefficient can be used as a measure of biomass inequality among individuals and has been previously used to assess temporal dynamics of seaweed populations structure (Bendel et al., 1989; Arenas & Fernández, 2000; Rivera & Scrosati, 2008). The coefficient ranges between 0 and 1, with 1 being the highest possible inequality (for instance: one individual account for all the biomass within an experimental unit).

### Dimensions of stability: resilience and recovery

We estimated the resilience and recovery of holdfast densities (juveniles and adults), percent cover of primary space, frond density, frond length, population biomass, and the Gini coefficient (population structure). Since recruitment was mostly seasonal, we did not estimate stability dimensions for this variable. The stability responses were estimated from log response ratios (LRR) calculated between disturbed and control plots as:

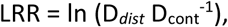

where D*_dist_* and D*_cont_* represent one of the measured attributes in the disturbed and control plots respectively. LRRs were estimated for pairs of control and disturbed plots, and for each sampling time. The pairing of plots was restricted to each block (two pairs per block).

Two dimensions of stability were calculated for each measured attribute. *Resilience (b)* was estimated as the slopes of regression of LRRs over time as:

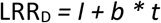

Where *I*, *b,* and *t* represent the intercept, slope, and time respectively. Values of *b* = 0 represent no change over time (lack of resilience) and values of *b* > 0 represents more rapid recovery over time (Donohue et al., 2016; Hillebrand et al., 2018; Radchuck et al., 2019). For the estimation of resilience, we considered LRR values calculated after the removal (from the second sampling time). *Recovery* (*c*) was estimated as the LRR calculated at the end of the experiment at each site (after 25 months in both sites). For all the population attributes, *c* = 0 represents maximum recovery, *c* < 0 denotes incomplete recovery, and *c* > 0 denotes overcompensation relative to control.

### Statistical analyses

Separate general linear models were used to assess the effect of experimental treatments on temporal patterns of *Durvillaea* recruitment, holdfast densities, percent cover, frond density, mean frond length, population biomass, and population structure (Gini coefficient). Model parameters were estimated through maximum likelihood, and we used a gaussian error distribution for all models. Model diagnostics were checked through visual inspection of quantile-quantile plots and fitted vs. adjusted residual plots.

One sample t-tests were used to assess if resilience and recovery of each population attribute were significantly different from zero (reference values). All the graphs and analyses were performed in R programming environment 4.0.2, using the “sjPlot” (Lüdecke, 2023), “ggplot2” (Wickham, 2016), “stats” (R Core Team, 2021) and “dineq” (Schulenberg, 2018) packages.

## RESULTS

### Temporal dynamics of population attributes

In Chaihuín, recruitment was not affected by the experimental removal and less than 20 individuals per m^2^ were recorded most of the time (Figure 1A). Two peaks of recruitment (i.e., up to 60 individuals per m^2^) occurred in both treatments in November of 2020 and in November of 2021 (Figure 1A; Table S1). Regardless of the experimental treatment, the highest densities of juveniles (i.e. > 20 ind. m^-2^) occurred in February of 2021 and February of 2022 (Figure 1B, Table S1). Statistically significant between-group differences in juvenile’ density were not found after the disturbance and across the experiment (Table S1). The density of adults decreased due the removal in March of 2020 to 1 ind. m^-2^ (Figure 1C; Table S1); then, similar densities in both treatments occurred after 7 months since the removal, in August of 2020 (Figure 1C). Two months after the removal (March of 2020), *Durvillaea* covered significatively less primary space in disturbed plots than in controls (Figure 1D; Table S1). During the following months, cover increased gradually until reaching statistically the same cover as controls in August of 2021, after 7 months since the removal (Figure 1D, Table S1).

**Figure 1.**
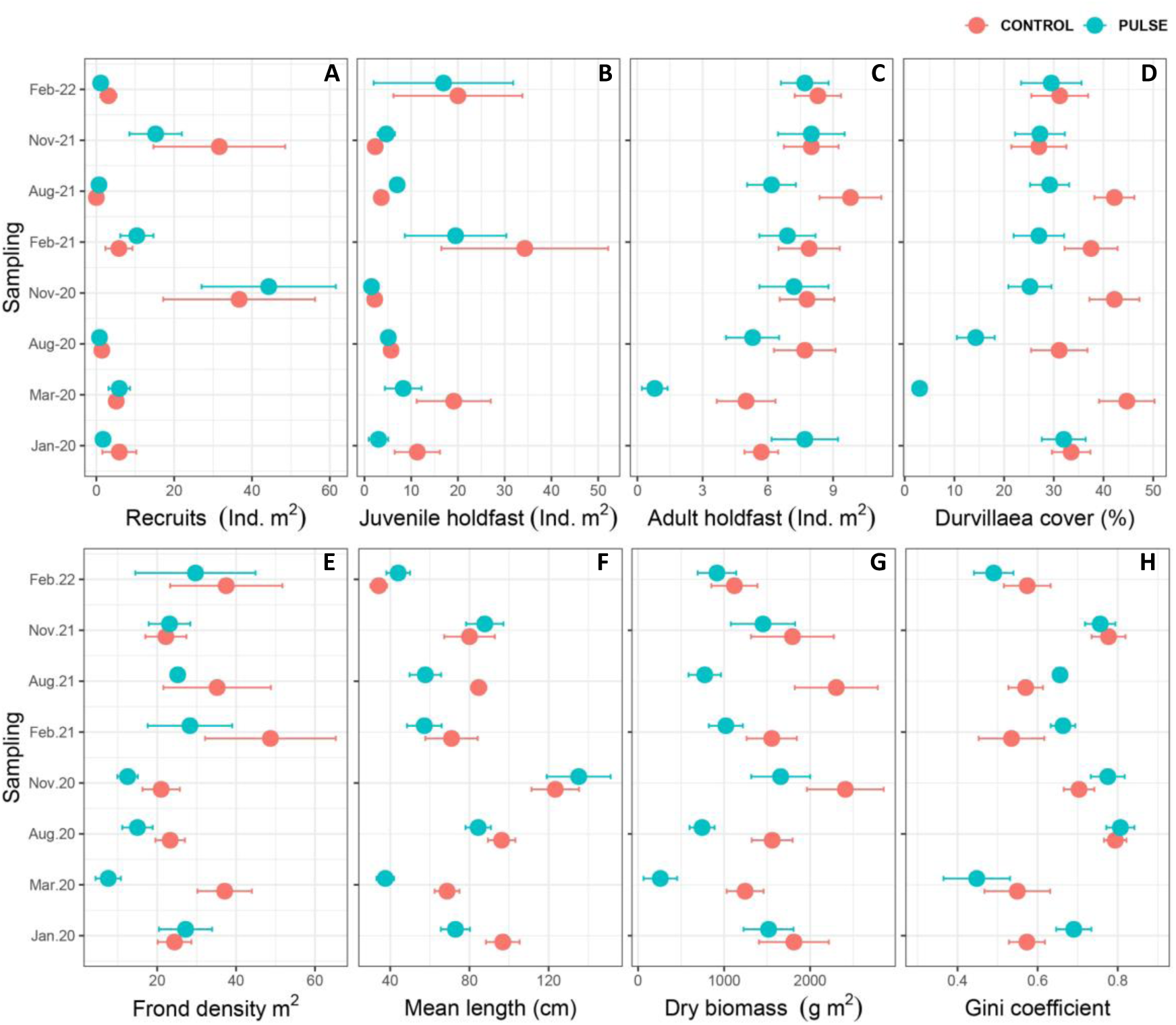
Temporal patterns of *Durvillaea incurvata* population from Chaihuín. We include density of recruits (A), juveniles (B) and adults (C) individuals, the total percent cover (D), frond density (E), mean frond length (F), population biomass (G) and population structure measured as the Gini coefficient of inequality (H) in control and pulse disturbed plots (red and blue dots respectively). The pulse disturbance was applied in January of 2020, immediately after first sampling. Horizontal bars represent ± standard error. Higher Gini coefficient values represent more inequality among fronds (i.e. the biomass is concentrated in some individuals within the population).

In Chaihuín and two months after the removal, we detected fewer fronds, smaller, and less than half of the biomass in disturbed plots than in controls (Figures 1E, F and G, Table S2). During the following months, frond density remained around 20 – 40 fronds m^2^ in both treatments, reaching higher densities in February of 2021 (Figure 1E, Table S2). The mean size of fronds ranged from 40 cm to 150 cm during the experiment; regardless of the experimental treatment, mean frond length was smallest in summer and largest during late winter and spring (Figure 1F; Table S2). In addition, population biomass in the disturbed plots remained several months around 200 – 300 g below the mean of controls (Figure 1G). However, statistically significant between-group differences were not detected for population biomass after the removal or thereafter (Table S2). The removal did not strongly impact the population structure in disturbed plots (Figure 1H; Table S2). Across the experiment, high values of the Gini coefficient indicated high population inequality in both treatments, because few large individuals accounted for most of the population biomass (Figure 1H).

In Loncoyén, recruits’ density ranged from 1 to more than 60 ind. m^-2^ during the experiment (Figure 2A). Mean recruit density did not differ between treatments after the experimental disturbance, or in the following sampling times (Table S3). For both treatments, densities of recruits peaked in October of 2020 and 2021 (Figure 2A). The density of juveniles was low, ranged from 1 to 30 individuals during the experiment, and were similar in both treatments after the disturbance (Figure 2B; Table S3). The density of adults differed between treatments two months after the disturbance (Figure 2C; Table S3) and took 5 months in disturbed plots to reach similar densities as controls (Figure 2C; Table S3). Two months after the experimental disturbance and in the following sampling times, *Durvillaea* covered less primary space in disturbed plots than in controls (Figure 2D; Table S3). Then, percent cover of *Durvillaea* increased steadily for 22 months until reach same cover as controls in August of 2022 (Figure 2D; Table S3).

**Figure 2.**
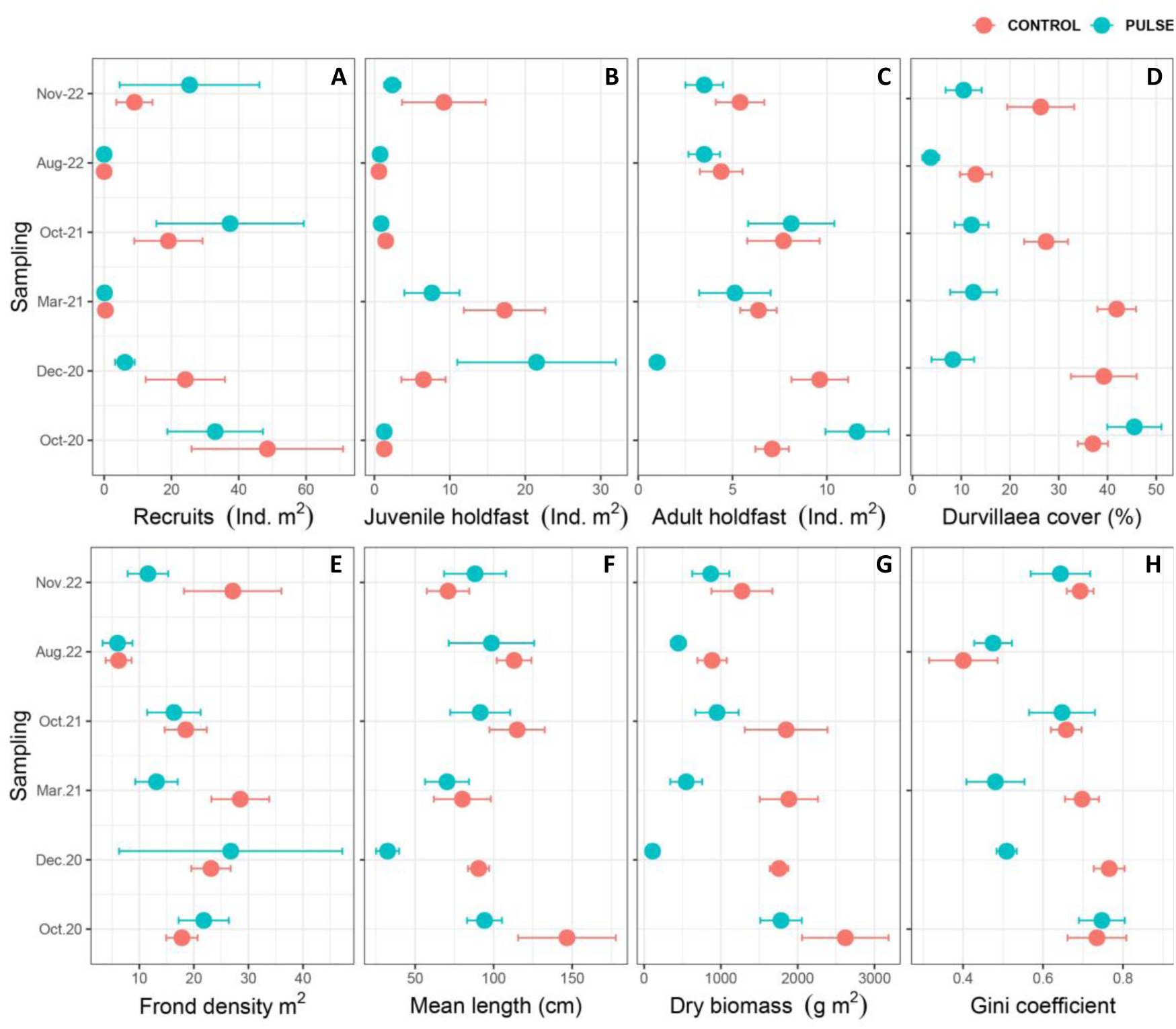
Temporal patterns of *Durvillaea incurvata* population from Loncoyén. We include density of recruits (A), juveniles (B) and adults (C) individuals, the total percent cover (D), frond density (E), mean frond length (F), population biomass (G) and population structure measured as the Gini coefficient of inequality (H) in control and pulse disturbed plots (red and blue dots respectively). The pulse disturbance was applied in October of 2020, immediately after first sampling. Horizontal bars represent ± standard error. For Gini coefficient higher values represent more inequality within fronds (i.e. the biomass is concentrated in some individuals within the population).

Mean frond density did not differ between control and disturbed plots after kelp removal in Loncoyén (December 2020; Figure 2E, Table S4). At this time, the mean frond length was near to 40 cm lower in disturbed than control plots (Figure 2F); however, statistically significant between-group differences were not found (Table S4). Similarly, less biomass was observed in disturbed than control plots (Figure 2G), but statistically significant between-group differences were not found at this time or thereafter (Table S4). After the removal, Gini values decreased respect to controls, indicating a homogenization of population size structure and the predominance of small size individuals accounting for most of population biomass (Figure 2H; Table S4). High inequality was common during most of the monitoring time (Figure 2H) and, regardless of the experimental treatment, inequality in frond biomass decreased in August of 2022 (Figure 2H; Table S4), probably associated to additional juvenile and adult mortality events produced by the strong wave action on this month.

### Population resilience and recovery

In Chaihuín, we detected a statistically significant and positive slope (i.e., “positive resilience”) of percent cover and frond density (Table 2). In contrast, the resilience of juvenile’s density, adults’ density, mean frond size, population structure, and biomass were so high (recovery occurred in less than 7 months), that resilience estimates of these attributes were not different from 0 (Table 2). The population from Loncoyén showed positive resilience for adults’ density, mean frond length, and biomass (Table 2), while the resilience of juvenile’s density, percent cover, frond density, and population structure resulted statistically equal to zero (Table 2). At the end of monitoring and after 25 months since the removal, *Durvillaea* populations from Chaihuín and Loncoyén showed complete recovery to the disturbance in all the measured attributes (mean value not different from 0; Table 2).

**Table 2.** Results of one sample t-test (mu = 0) for resilience and recovery of the five attributes measured in *Durvillaea incurvata* populations in two intertidal rocky sites (Chaihuín and Loncoyén). For all measured responses, high, low and no resilience corresponds to estimates values of > 0, < 0 and = 0, respectively. While recovery is high for estimates values = 0, and low when values are higher or lower than 0.

|  |  | Chaihuín |  |  |  |  |  | Loncoyén |  |  |  |  |  |
| --- | --- | --- | --- | --- | --- | --- | --- | --- | --- | --- | --- | --- | --- |
| Dimension | Response | Estimate | t | df | p-value | CI LOW | CI HIGH | Estimate | t | df | p-value | CI LOW | CI HIGH |
| Resilience | Juvenile holdfast density | 0.044 | 0.178 | 9 | 0.862 | -0.513 | 0.600 | -0.280 | -0.552 | 7 | 0.598 | -1.480 | 0.918 |
|  | Adult holdfast density | 0.128 | 1.540 | 9 | 0.157 | -0.059 | 0.315 | 0.367 | 2.540 | 9 | **0.032** | 0.039 | 0.694 |
|  | Cover | **0.373** | **4.910** | **9** | **<0.001** | 0.201 | 0.545 | 0.321 | 1.660 | 9 | 0.131 | -0.116 | 0.757 |
|  | Fronde density | **0.268** | **3.605** | **9** | **0.005** | 0.099 | 0.436 | 0.187 | 0.833 | 7 | 0.432 | -0.344 | 0.719 |
|  | Fronde length | 0.042 | 0.511 | 9 | 0.622 | -0.143 | 0.226 | 0.421 | 2.620 | 9 | **0.028** | 0.057 | 0.785 |
|  | Biomass | 0.338 | 1.850 | 9 | 0.097 | -0.075 | 0.751 | 0.661 | 3.000 | 9 | **0.015** | 0.162 | 1.160 |
|  | Structure | -0.022 | -0.844 | 9 | 0.420 | -0.083 | 0.038 | 0.171 | 1.82 | 3 | 0.166 | -0.128 | 0.470 |
| Recovery | Juvenile holdfast density | -0.343 | -0.388 | 9 | 0.707 | -2.340 | 1.660 | -1.070 | -1.660 | 5 | 0.157 | -2.720 | 0.582 |
|  | Adult holdfast density | -0.140 | -0.800 | 9 | 0.444 | -0.534 | 0.255 | -0.579 | -1.320 | 8 | 0.222 | -1.590 | 0.429 |
|  | Cover | -0.011 | -0.026 | 9 | 0.980 | -0.956 | 0.935 | -1.090 | -1.860 | 9 | 0.096 | -2.400 | 0.233 |
|  | Fronde density | -0.256 | -0.438 | 9 | 0.671 | -1.578 | 1.066 | -0.871 | -1.274 | 7 | 0.243 | -2.488 | 0.746 |
|  | Fronde length | 0.199 | 0.932 | 9 | 0.376 | -0.285 | 0.684 | -0.783 | -1.090 | 8 | 0.306 | -2.440 | 0.869 |
|  | Biomass | -0.091 | -0.271 | 9 | 0.792 | -0.847 | 0.667 | -1.900 | -1.720 | 9 | 0.119 | -4.390 | 0.597 |
|  | Structure | -0.192 | -0.769 | 9 | 0.461 | -0.759 | 0.373 | -0.071 | -0.451 | 3 | 0.683 | -0.569 | 0.423 |

## DISCUSSION

After an experimental removal of *Durvillaea incurvata* in south – central Chile, which mimicked the way that gatherers harvest this species at small spatial scales, we observed high resilience and complete recovery of holdfast densities, percent cover, frond density, frond mean size, population biomass, and population structure. In two intertidal sites, Chaihuín and Loncoyén, these attributes converged toward the reference values and recovered relatively fast (within five to seven months), except for the percent cover of *Durvillaea* in Loncoyén, which took almost two years to recover. In addition, recruitment was unaffected by the experimental removal and did not follow the patterns of adult abundances in experimental plots. Therefore, for both studied populations, our evidence supports the hypothesis that constant (rather than massive) recruitment and high growth rates allow for rapid recovery (i.e. within a year) of *Durvillaea* populations after disturbances. The strong stability responses of these populations suggest that harvesting at small spatial scale, within dense kelp stands and removing the whole individuals (including the holdfast, stipes, and fronds), has the potential to sustain harvesting by local gatherers without long-lasting impacts on the kelp populations.

### Population dynamics after disturbances

Surprisingly, the removal of *Durvillaea* did not have a significant impact on recruitment, which seemed to be more linked to environmental seasonal variability. Both populations experienced substantial recruitment events during the spring, sometimes reaching densities of approximately 200 individuals m^-2^. Due the short lifespan of *Durvillaea* zygotes, which settle within a few hours and within a short distance from their progenitors (Dunmore, 2006), we suggest that inputs of recruits came from adult individuals observed near to the experimental plots (see Schiel & Foster 2006 for an example elsewhere). Our results are consistent with the strong responses exhibited by the intertidal laminarian kelp *Lessonia berteorana* in northern Chile, whose rapid colonization and high growth rates boosted population resilience and recovery in less than 10 months (Westermeier et al. 2019). Similarly, after experimental removal, enhanced light availability and empty substrate allowed high recolonization and promoted the recovery of intertidal and subtidal laminarian kelps such as *Lessonia* spp. (Ojeda & Santelices, 1984; Vásquez et al. 2012, Vega et al. 2014; Gouraguine et al. 2021,) and fucoids such as *Fucus* sp, *Durvillaea* spp. and *Hormosira banksii* (Sandone, 1991; Taylor & Schiel, 2005; Bellgrove et al., 2010). Therefore, we suggest that after our experimental removal of *Durvillaea*, recovery was enhanced by high propagule availability provided by the surrounding undisturbed population and by the utilization of an enhanced light environment in absence of adults.

Early life stages of fucoid and laminarian kelps may have important implications for population recovery after disturbances (Schiel & Foster, 2006). After settlement, for example, the gametophytes of laminarian kelps such as *Macrocystis pyrifera* can delay their development and form a bank of microscopic forms that persist under unfavorable conditions (Yoneshigue-Valentin, 1990; Dieck, 1993; Schiel & Foster, 2006). Similarly, the microscopic zygotes of *Fucus distichus* and *Pelvetia fastigiata* can take up to several months to reach macroscopic forms and growth may be delayed when conditions are unfavorable (Gunnil, 1980; Ang 1991). Similarly, after the canopy removal of *Ecklonia radiata*, pre-existing juveniles can survive to post-disturbance conditions and contribute to a rapid recovery of the population (Toohey et al., 2007). Thus, along with high propagule availability, it is likely that a set of already settled individuals took advantage of the enhanced light environment after the removal of *Durvillaea* and contributed to boost resilience and a rapid population recovery.

Despite that the survival of early life stages of fucoids and laminarian kelps are mostly impacted by density-dependent processes (Gunnil, 1980; Schiel & Foster, 2006), in *Durvillaea* spp. the coalesce among individuals during growth constitutes a mechanism by which population density increases and intraspecific competition for primary space is alleviated (Ojeda & Santelices, 1984). Coalescent individuals can reach higher survival rates than isolated individuals, which can provide with stability to the population (Bustamante & Castilla, 1989; Westermeier et al., 1994; Wernberg, 2005; Santelices & Aedo, 2006; Santelices & Alvarado, 2008). In addition, large holdfasts (i.e. > 10 cm in diameter) usually support numerous stipes and fronds, sustaining higher amounts of biomass than single individuals (Santelices, 1980; Santelices et al., 1980; Ojeda & Santelices, 1984; Velásquez et al., 2020). In our monitored sites, fewer adult coalescent individuals were reported in disturbed plots after the removal and in the following months (Supplementary figure 1). Therefore, recovery rates of attributes such as frond density, percent cover, and population biomass depend on the rate of coalescence.

### High resilience and recovery of Durvillaea

The stability responses observed in our study were faster than those previously reported for *Durvillaea* populations and for some fucoid and laminarian kelps across world temperate reefs (Westermeier et al. 1994; Levitt et al. 2002, Castilla et al., 2007; Tait & Schiel, 2011; Wernberg et al. 2019; Wernberg et al., 2020; Thomsen et al. 2021). Several factors may explain this discrepancy. For example, pruning the kelp fronds near to the holdfast (instead of removing whole individuals as in our study), could reduce the ability to regenerate fronds, promote the recruitment on holdfasts that later detach from primary substrate, resulting in delayed or lack of recovery in populations of *D. antarctica* (Westermeier et al. 1994), *Ecklonia maxima* (Levitt et al. 2002), and *L. berteorana* (Vásquez et al., 2012). Also, interspecific interactions can be critical for population recovery after disturbances (Wernberg et al., 2019). For example, the rapid establishment of non-native kelps, opportunistic species such as *Ulva* spp., crustose seaweeds, and turf forming seaweeds, may delay and inhibit the recovery of kelp populations after disturbances through habitat modification (Schiel et al., 2018; Thomsen et al., 2019, 2021), competition for primary substrata, and reduction of propagule pressure (Jenkins et al., 2004; Petraitis & Dudgeon, 2004; Smale, 2020). Kelps such as *L. spicata* and *M. pyrifera*, which can delay the recovery of *Durvillaea* populations (Santelices, 1990; Santelices et al., 1980; Westermeier et al., 1994; Moreno, 2001), were almost absent during our experiment. *Lessonia spicata* occurred at low densities in Loncoyén, while *M. pyrifera* was common in both sites but sporophytes were not longer than 40 cm. Then, alternative kelps were unable to avoid the high recruitment and growth rate of *Durvillaea* after the removal. Finally, mass mortality of kelps as consequence of large-scale disturbances such as earthquake-induced coastal uplift, prolonged exposure to marine heat waves, and intense harvesting can reduce population recovery and even cause the local extinction of some species (Castilla et al., 2007; Wernberg et al., 2019; Thomsen et al., 2021). These events drastically drop the amount of parental stock within disturbed areas and can reduce the resilience and recovery of densities, percent cover, biomass, and productivity of fucoid and laminarian kelp populations such as *L. trabeculata* (Bularz et al., 2022), *Durvillaea* spp. in central Chile and in New Zealand (Castilla et al., 2007; Tait & Schiel 2011; Schiel et al., 2018; Thomsen et al., 2019, 2021), *Saccharina latissima* in the North Atlantic (Filbee-Dexter et al., 2020), and several kelp populations across Australia (Layton et al., 2019; Wernberg et al., 2019; 2020). Therefore, the small-scale impact of our experimental removal within dense stands of adults, seems to be key to sustain the self-replacement of *Durvillaea* after disturbances, allowing a high resilience and recovery of the population within a short period of time.

### Conclusions

In summary, this study suggest that bull kelp can recover from small scale disturbances, such as the removal of whole individuals within 1 m^2^ plots distributed among dense kelp stands. This kind of harvesting is common in southern Chile and seems to be beneficial to ensure population self-replacement, by leaving the presence of parental stock within harvesting zones. Furthermore, the continuous recruitment within available space, along with high growth rates of fronds where key to enhance recovery rates of the monitored populations after the disturbance. In addition, the timing of the disturbance may drive different stability responses when done in different seasons. Accordingly, future management strategies of commercially and ecologically important intertidal kelps in southern Chile and elsewhere should prioritize small-scale harvesting during the periods of higher recruitment rates to promote a rapid recovery and ensure the long-term sustainability and stability of these valuable species and the associated benefits provided to coastal communities.

