## Supplementary figure and tables for "Rapid population recovery of a foundation species from experimental small-scale harvesting"

### Supplementary material

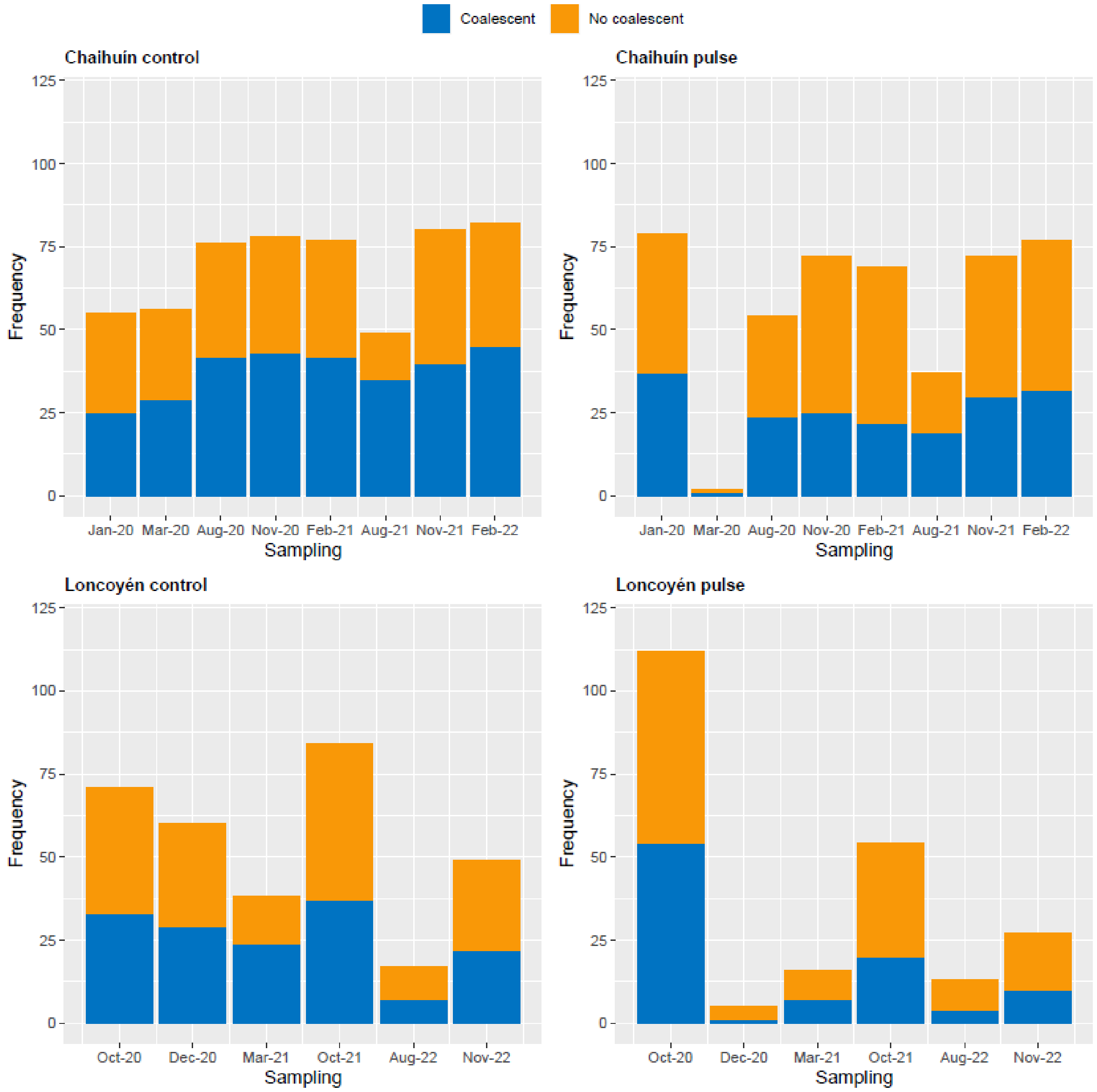


**Figure S1.** Frequency of adult (> 5 cm holdfast diameter) coalescent and non-coalescent individuals, found in ten control and ten pulse disturbed plots in Chaihuín and Loncoyén.

**Table S1.** General linear model results for population traits densities and cover of *Durvillaea incurvata* in Chaihuín (southern Chile). Intercept represents January 2020 in the control.

| | Recruits density | | | Juveniles density | | | Adults density | | | Percent cover | | |
|---|---|---|---|---|---|---|---|---|---|---|---|---|
| *Predictors* | *Estimates* | *Std. Error* | *p* | *Estimates* | *Std. Error* | *p* | *Estimates* | *Std. Error* | *p* | *Estimates* | *Std. Error* | *p* |
| Intercept | 5.90 | 8.48 | 0.488 | 11.30 | 8.05 | 0.163 | 5.70 | 1.27 | **<0.001** | 33.50 | 4.85 | **<0.001** |
| Treatment (T) | -4.20 | 12.00 | 0.727 | -8.30 | 11.38 | 0.467 | 2.00 | 1.79 | 0.267 | -1.50 | 6.86 | 0.827 |
| Mar-20 | -0.80 | 12.00 | 0.947 | 7.80 | 11.38 | 0.494 | -0.70 | 1.79 | 0.697 | 11.20 | 6.86 | 0.105 |
| Aug-20 | -4.50 | 12.00 | 0.708 | -5.60 | 11.38 | 0.624 | 2.00 | 1.79 | 0.267 | -2.40 | 6.86 | 0.727 |
| Nov-20 | 30.80 | 12.00 | **0.011** | -9.10 | 11.38 | 0.425 | 2.10 | 1.79 | 0.244 | 8.70 | 6.86 | 0.207 |
| Feb-21 | -0.10 | 12.00 | 0.993 | 23.00 | 11.38 | **0.045** | 2.20 | 1.79 | 0.222 | 4.00 | 6.86 | 0.561 |
| Aug-21 | -5.90 | 14.70 | 0.689 | -7.70 | 13.94 | 0.582 | 4.10 | 2.20 | 0.064 | 8.70 | 8.40 | 0.302 |
| Nov-21 | 25.70 | 12.00 | **0.034** | -9.00 | 11.38 | 0.430 | 2.30 | 1.79 | 0.202 | -6.50 | 6.86 | 0.345 |
| Feb-22 | -2.80 | 12.00 | 0.816 | 8.70 | 11.38 | 0.446 | 2.60 | 1.79 | 0.150 | -2.30 | 6.86 | 0.738 |
| T:Mar-20 | 5.00 | 16.97 | 0.769 | -2.50 | 16.10 | 0.877 | -6.20 | 2.54 | **0.016** | -40.20 | 9.69 | **<0.001** |
| T:Aug-20 | 3.60 | 16.97 | 0.832 | 7.70 | 16.10 | 0.633 | -4.40 | 2.54 | 0.085 | -15.30 | 9.69 | 0.117 |
| T:Nov-20 | 11.80 | 16.97 | 0.488 | 7.60 | 16.10 | 0.638 | -2.60 | 2.54 | 0.307 | -15.50 | 9.69 | 0.112 |
| T:Feb-21 | 8.80 | 16.97 | 0.605 | -6.50 | 16.10 | 0.687 | -3.00 | 2.54 | 0.239 | -9.00 | 9.69 | 0.355 |
| T:Aug-21 | 4.87 | 20.20 | 0.810 | 11.70 | 19.16 | 0.542 | -5.63 | 3.02 | 0.064 | -11.53 | 11.54 | 0.319 |
| T:Nov-21 | -12.18 | 17.20 | 0.480 | 10.67 | 16.32 | 0.514 | -2.00 | 2.57 | 0.438 | 1.72 | 9.83 | 0.861 |
| T:Feb-22 | 2.20 | 16.97 | 0.897 | 5.20 | 16.10 | 0.747 | -2.60 | 2.54 | 0.307 | -0.20 | 9.69 | 0.984 |
| Observations | 150 | | | 150 | | | 150 | | | 150 | | |
| $R^2$ | 0.230 | | | 0.125 | | | 0.210 | | | 0.328 | | |

**Table S2.** General linear model results for population traits of *Durvillaea incurvata* in Chaihuín (southern Chile). Intercept represents January 2020 in the control.

| | **Frond density** | | | **Mean length** | | | **Dry biomass** | | | **Size structure** | | |
|---|---|---|---|---|---|---|---|---|---|---|---|---|
| *Predictors* | *Estimates* | *Std. Error* | *p* | *Estimates* | *Std. Error* | *p* | *Estimates* | *Std. Error* | *p* | *Estimates* | *Std. Error* | *p* |
| Intercept | 24.40 | 8.62 | **0.005** | 96.86 | 8.79 | **<0.001** | 1811.05 | 312.31 | **<0.001** | 0.57 | 0.05 | **<0.001** |
| Treatment (T) | 2.80 | 12.19 | 0.819 | -23.94 | 12.43 | 0.056 | -294.89 | 441.67 | 0.505 | 0.12 | 0.07 | 0.076 |
| Mar-20 | 12.70 | 12.19 | 0.299 | -28.17 | 12.43 | **0.025** | -567.32 | 441.67 | 0.201 | -0.02 | 0.07 | 0.760 |
| Aug-20 | -1.10 | 12.19 | 0.928 | -0.58 | 12.43 | 0.963 | -252.72 | 441.67 | 0.568 | 0.22 | 0.07 | **<0.001** |
| Nov-20 | -3.40 | 12.19 | 0.781 | 26.51 | 12.43 | **0.035** | 597.89 | 441.67 | 0.178 | 0.13 | 0.07 | **0.042** |
| Feb-21 | 24.30 | 12.19 | **0.048** | -25.91 | 12.43 | **0.039** | -257.87 | 441.67 | 0.560 | -0.04 | 0.07 | 0.623 |
| Aug-21 | 10.80 | 14.93 | 0.471 | -12.18 | 15.23 | 0.425 | 491.86 | 540.94 | 0.365 | -0.00 | 0.09 | 0.968 |
| Nov-21 | -2.20 | 12.19 | 0.857 | -16.82 | 12.43 | 0.178 | -17.62 | 441.67 | 0.968 | 0.20 | 0.07 | **0.002** |
| Feb-22 | 13.10 | 12.19 | 0.285 | -62.68 | 12.43 | **<0.001** | -691.96 | 441.67 | 0.120 | 0.00 | 0.07 | 0.990 |
| T:Mar-20 | -32.30 | 17.24 | 0.063 | -7.26 | 17.58 | 0.680 | -690.80 | 624.62 | 0.271 | -0.22 | 0.11 | 0.060 |
| T:Aug-20 | -11.10 | 17.24 | 0.521 | 12.09 | 17.58 | 0.493 | -520.24 | 624.62 | 0.406 | -0.10 | 0.10 | 0.227 |
| T:Nov-20 | -11.30 | 17.24 | 0.513 | 35.78 | 17.58 | **0.044** | -457.35 | 624.62 | 0.465 | -0.05 | 0.10 | 0.619 |
| T:Feb-21 | -23.20 | 17.24 | 0.181 | 10.21 | 17.58 | 0.563 | -238.37 | 624.62 | 0.703 | 0.01 | 0.10 | 0.906 |
| T:Aug-21 | -12.80 | 21.11 | 0.545 | -3.01 | 21.54 | 0.889 | -1233.87 | 743.44 | 0.099 | -0.03 | 0.13 | 0.758 |
| T:Nov-21 | -1.87 | 17.77 | 0.916 | 31.66 | 18.13 | 0.083 | -47.17 | 633.23 | 0.941 | -0.14 | 0.11 | 0.140 |
| T:Feb-22 | -10.60 | 17.24 | 0.540 | 33.76 | 17.58 | 0.057 | 92.10 | 624.62 | 0.883 | -0.20 | 0.10 | **0.041** |
| Observations | 148 | | | 148 | | | 150 | | | 146 | | |
| $R^2$ | 0.135 | | | 0.534 | | | 0.248 | | | 0.378 | | |

**Table S3.** General linear model results for population densities and cover of *Durvillaea incurvata* in Loncoyén (southern Chile). Intercept represents October of 2020 in the control.

| | **Recruits density** | | | **Juveniles density** | | | **Adults density** | | | **Percent cover** | | |
|---|---|---|---|---|---|---|---|---|---|---|---|---|
| *Predictors* | *Estimates* | *Std. Error* | *p* | *Estimates* | *Std. Error* | *p* | *Estimates* | *Std. Error* | *p* | *Estimates* | *Std. Error* | *p* |
| Intercept | 48.50 | 13.56 | **0.001** | 1.30 | 3.80 | 0.733 | 7.10 | 1.48 | **<0.001** | 37.00 | 4.74 | **<0.001** |
| Treatment (T) | -15.50 | 19.18 | 0.421 | 0.00 | 5.37 | 1.000 | 4.50 | 2.10 | **0.035** | 8.50 | 6.71 | 0.209 |
| Dec-20 | -24.38 | 20.34 | 0.234 | 5.20 | 5.69 | 0.364 | 2.53 | 2.22 | 0.260 | 2.25 | 7.12 | 0.753 |
| Mar-21 | -48.13 | 20.34 | **0.020** | 15.95 | 5.69 | **0.006** | -0.72 | 2.22 | 0.745 | 4.87 | 7.12 | 0.495 |
| Oct-21 | -29.40 | 19.18 | 0.129 | 0.20 | 5.37 | 0.970 | 0.60 | 2.10 | 0.776 | -9.60 | 6.71 | 0.156 |
| Aug-22 | -48.50 | 23.49 | **0.042** | -0.70 | 6.57 | 0.915 | -2.70 | 2.57 | 0.296 | -24.00 | 8.22 | **0.004** |
| Nov-22 | -39.50 | 19.18 | **0.043** | 7.90 | 5.37 | 0.145 | -1.70 | 2.10 | 0.420 | -10.70 | 6.71 | 0.115 |
| T:Dec-20 | -2.46 | 30.07 | 0.935 | 15.00 | 8.42 | 0.078 | -13.13 | 3.29 | **<0.001** | -39.46 | 10.26 | **<0.001** |
| T:Mar-21 | 15.25 | 28.77 | 0.597 | -9.62 | 8.05 | 0.235 | -5.75 | 3.15 | 0.071 | -37.87 | 10.07 | **<0.001** |
| T:Oct-21 | 33.84 | 27.50 | 0.222 | -0.61 | 7.70 | 0.937 | -4.09 | 3.01 | 0.178 | -23.79 | 9.62 | **0.015** |
| T:Aug-22 | 15.50 | 34.58 | 0.655 | 0.15 | 9.68 | 0.988 | -5.40 | 3.78 | 0.157 | -17.75 | 12.10 | 0.146 |
| T:Nov-22 | 31.88 | 27.96 | 0.257 | -6.83 | 7.83 | 0.386 | -6.40 | 3.06 | **0.039** | -24.30 | 9.78 | **0.015** |
| Observations | 96 | | | 96 | | | 96 | | | 97 | | |
| $R^2$ | 0.139 | | | 0.239 | | | 0.280 | | | 0.492 | | |

**Table S4.** General linear model results for population traits of *Durvillaea incurvata* in Chaihuín Loncoyén (southern Chile). Intercept represents October of 2020 in the control.

| | **Frond density** | | | **Mean length** | | | **Dry biomass** | | | **Size structure** | | |
|---|---|---|---|---|---|---|---|---|---|---|---|---|
| *Predictors* | *Estimates* | *Std. Error* | *p* | *Estimates* | *Std. Error* | *p* | *Estimates* | *Std. Error* | *p* | *Estimates* | *Std. Error* | *p* |
| Intercept | 17.80 | 5.02 | **0.001** | 146.68 | 15.60 | **<0.001** | 2620.22 | 348.15 | **<0.001** | 0.73 | 0.06 | **<0.001** |
| Treatment (T) | 4.00 | 7.10 | 0.575 | -52.50 | 22.06 | **0.020** | -836.92 | 492.36 | 0.093 | 0.01 | 0.08 | 0.884 |
| Dec-20 | 5.33 | 7.54 | 0.482 | -56.27 | 23.39 | **0.019** | -862.24 | 522.23 | 0.102 | 0.03 | 0.09 | 0.711 |
| Mar-21 | 10.70 | 7.54 | 0.160 | -66.61 | 23.39 | **0.006** | -735.38 | 522.23 | 0.163 | -0.04 | 0.09 | 0.659 |
| Oct-21 | 0.70 | 7.10 | 0.922 | -31.73 | 22.66 | 0.165 | -770.01 | 492.36 | 0.122 | -0.08 | 0.08 | 0.334 |
| Aug-22 | -11.60 | 8.70 | 0.186 | -33.69 | 27.01 | 0.216 | -1734.67 | 603.01 | **0.005** | -0.33 | 0.12 | **0.005** |
| Nov-22 | 9.31 | 7.30 | 0.206 | -75.72 | 22.66 | **0.001** | -1346.12 | 492.36 | **0.008** | -0.04 | 0.08 | 0.603 |
| T:Dec-20 | -0.38 | 12.05 | 0.975 | -5.45 | 37.40 | 0.884 | -810.20 | 771.97 | 0.297 | -0.27 | 0.12 | **0.023** |
| T:Mar-21 | -19.36 | 10.87 | 0.079 | 42.73 | 33.73 | 0.209 | -499.21 | 738.54 | 0.501 | -0.23 | 0.13 | 0.075 |
| T:Oct-21 | -6.17 | 10.85 | 0.572 | 28.98 | 33.23 | 0.386 | -63.24 | 705.90 | 0.929 | -0.02 | 0.12 | 0.854 |
| T:Aug-22 | -4.20 | 12.81 | 0.744 | 38.11 | 39.76 | 0.341 | 398.16 | 887.61 | 0.655 | 0.06 | 0.15 | 0.686 |
| T:Nov-22 | -19.54 | 10.70 | 0.072 | 69.68 | 33.23 | **0.039** | 431.24 | 717.73 | 0.550 | -0.06 | 0.14 | 0.647 |
| Observations | 88 | | | 88 | | | 96 | | | 87 | | |
| $R^2$ | 0.171 | | | 0.243 | | | 0.312 | | | 0.305 | | |
